# Omicron evolution drives increased ciliated cell tropism and dysfunction in nasal epithelia

**DOI:** 10.1101/2025.11.19.689397

**Authors:** Helen Mostafavi, Rhys H. Parry, Georgina McCallum, Julian D.J. Sng, Merja Joensuu, Kirsty R. Short, Andrii Slonchak, Alexander A. Khromykh

## Abstract

SARS-CoV-2 Omicron variants demonstrate greater replication efficiency in the upper respiratory tract than both ancestral virus and Delta variant, potentially due to stronger interactions with motile cilia in human nasal epithelial cells. Despite the emergence of numerous Omicron variants, there has been no comprehensive investigation comparing their interactions with ciliated cells. Here, we use differentiated primary human nasal epithelial cell (NEC) cultures to compare an ancestral isolate (QLD02) to Omicron variants BA.1, BA.5 and XBB. Transcriptomic analysis revealed that BA.5 and XBB infection, unlike BA.1 and QLD02, drives a distinct host response, defined by the profound downregulation of genes essential for cilium assembly and function at 48 hours post-infection. The BA.5/XBB signature was also characterized by the strong upregulation of pro-apoptotic and inflammatory pathways. Immunofluorescence of nucleocapsid-positive cells in NECs showed a significant 13-fold increase for BA.5 relative to QLD02. Additionally, BA.1 and BA.5 exhibited greater tropism for ciliated cells (7.6-fold for BA.1, 9-fold for BA.5) than QLD02. BA.5 and BA.1 infection were associated with elevated apoptosis in both infected and bystander cells, however, the total number of ciliated cells remained unchanged. Consistent with this, BA.5 infection significantly reduced DNAH5 outer dynein arm protein in the cilia, indicating dysfunction of cilia motility rather than overt cilia loss. Overall, we demonstrate a stepwise evolution at the nasal barrier: while enhanced ciliated cell tropism and elevated apoptosis are conserved features of Omicron infection, ciliary dysfunction, evident at both the transcriptional and protein level, is a more pronounced phenotype emerging with BA.5 and preserved in the XBB variant.

**Graphical Abstract:** 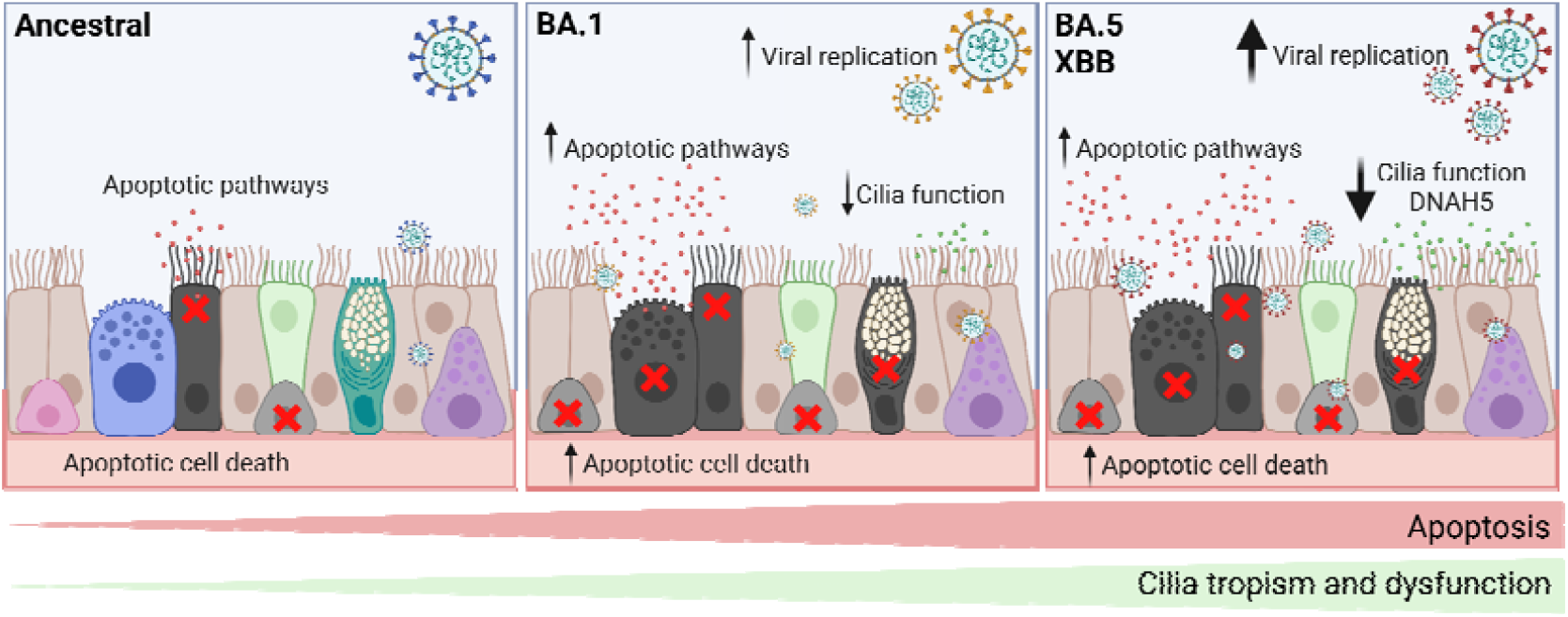

## Introduction

The upper respiratory tract is the primary site of infection with SARS-CoV-2 (1). The nasal mucosa is populated by a variety of different cells, including goblet cells, ciliated cells, mucus-producing cells, and nasal progenitor cells (2). *In vivo*, SARS-CoV-2 displays preferential tropism for multiciliated cells of the nasal epithelium (3, 4). For example, in nasal tissue of patients hospitalised in 2020 for COVID-19, SARS-CoV-2 nucleocapsid protein (NP) expression was restricted to multiciliated cells displaying acetylated α-tubulin and absent in goblet cells, secretory, or differentiating cells (5). Similarly, in *in vitro* air-liquid interface cultures of human nasal epithelial cells there were significantly increased levels of SARS-CoV-2 (ancestral and alpha variant) infection in ciliated cells and secretory-ciliated cells (4). In human challenge studies with the wildtype ancestral virus of SARS-CoV-2, single cell sequencing of nasopharyngeal swabs identified evidence of productive viral infection of both ciliated and goblet cells (6). Additionally, this study identified a subset of ciliated cells that, whilst only constituting 4% of total infected cells, contained 67% of detected viral RNA (6). Importantly, ancestral SARS-CoV-2 has itself been shown to downregulate key ciliogenesis regulators, including FOXJ1 and RFX3, as well as ciliary structural genes such as DNAH7, by 4 days post-infection (7). This establishes that cilia-disruptive transcriptional changes are not unique to Omicron variants, but rather that the extent and coordination of this response may have evolved with successive lineages.

Targeting ciliated cells has important implications for viral pathogenesis. Disruption and shedding of ciliary axonemes has been indicated to impair mucociliary clearance (8). This impaired mucociliary clearance likely facilitates the dissemination of virus to the lower respiratory tract, which may have implications for disease severity (7, 9). Disease severity may also be exacerbated by changes to cytokine production from infected and dying ciliated cells, resulting in a dysregulated immune response (8).

Ciliary motility is driven by the outer dynein arm (ODA) motor complex, of which DNAH5 is a principal heavy-chain component required for effective axonemal force generation. In humans, loss of DNAH5 is a canonical cause of primary ciliary dyskinesia, in which cilia may remain structurally present yet beat ineffectively or become dyskinetic, resulting in impaired mucociliary clearance (10-12). DNAH5 localisation at the ciliary axoneme therefore provides a biologically informative readout of ciliary motor competence that is distinct from simply quantifying cilia abundance. Whether SARS-CoV-2 disrupts ODA components such as DNAH5 in infected nasal epithelia has not been investigated (7, 8, 13).

Recent evidence suggests that the extent to which SARS-CoV-2 targets ciliated epithelial cells has evolved over time (14). The Omicron variant, which emerged in late 2021, exhibits enhanced replication efficiency in the upper respiratory tract compared to the ancestral virus and the Delta variant. This increased upper respiratory tropism is thought to be associated with increased transmissibility in Omicron variants (15). Consistent with these data, compared to the ancestral virus and Delta, Omicron was more likely to bind motile cilia in human nasal epithelial cells (13, 16).

Since its emergence in late 2021, Omicron has diversified substantially into a variety of different variants. The BA.1 and BA.2 variants diverged first from a common Omicron ancestor (B.1.1.529) in November of 2021. Subsequently, the BA.2 variant diverged again in early 2022, giving rise to the BA.4 and BA.5 variants, characterised by increased fusogenicity and antibody evasion, likely conferred by additional spike mutations such as L452R and F486V, respectively (17). By mid-2022, BA.5 had become the dominant global variant, and it remained as such until approximately December 2022. Recombination events between two BA.2 variants later gave rise to XBB, which subsequently became the predominant variant in several regions by early 2023. These different Omicron variants display marked differences in tropism and pathogenesis. For example, in human nasal epithelial cells XBB.1.5 replicated more efficiently than BA.2 (18). Whether this reflects increased targeting of ciliated cells in the upper respiratory tract remains unclear, although early studies from a limited number of nasal swabs (n = 4) suggested that the later Omicron variant XBC was more effective than BA.5 at targeting ciliated cells in the upper respiratory tract (19). Here, we comprehensively examine the effect of different Omicron variants on cilia in the upper respiratory tract, to better understand variant-dependent differences in pathogenesis.

## Results

### BA.1 and BA.5 Omicron variants of SARS-CoV-2 demonstrate higher replication efficiency compared to the ancestral virus in primary human nasal epithelial cells

To investigate different Omicron variants at the primary site of infection, we compared the replication of three key SARS-CoV-2 isolates in primary human nasal epithelial cells (NECs) (Fig. 1A). The selected variants, as indicated by whole-genome phylogeny (Fig. 1B), include: an ancestral (Pango lineage A) isolate (QLD02, Jan 2020), an early Omicron (Pango lineage BA.1) isolate (QLD2639C, Dec 2021), and a later Omicron (Pango lineage BA.5) variant isolate (QLD-QIMR03, Jul 2022). At 48 hours post-infection, intracellular viral RNA loads for four viral genes (E, S, RdRP[nsp12], and N) were quantified by RT-qPCR (Fig. 1C). Both BA.1 and BA.5 demonstrated higher viral RNA loads than the ancestral QLD02 virus. This enhanced replication was statistically significant for the S gene in both BA.1 (p = 0.0284) and BA.5 (p = 0.0133) compared to QLD02, and for the E gene, where BA.5 replication was significantly higher than ancestral (p = 0.0339), suggesting a progressive increase in intracellular replication efficiency in NECs with Omicron variants. There were no significant differences in RNA levels for any of the four genes between BA.1 and BA.5.

**Figure 1.**
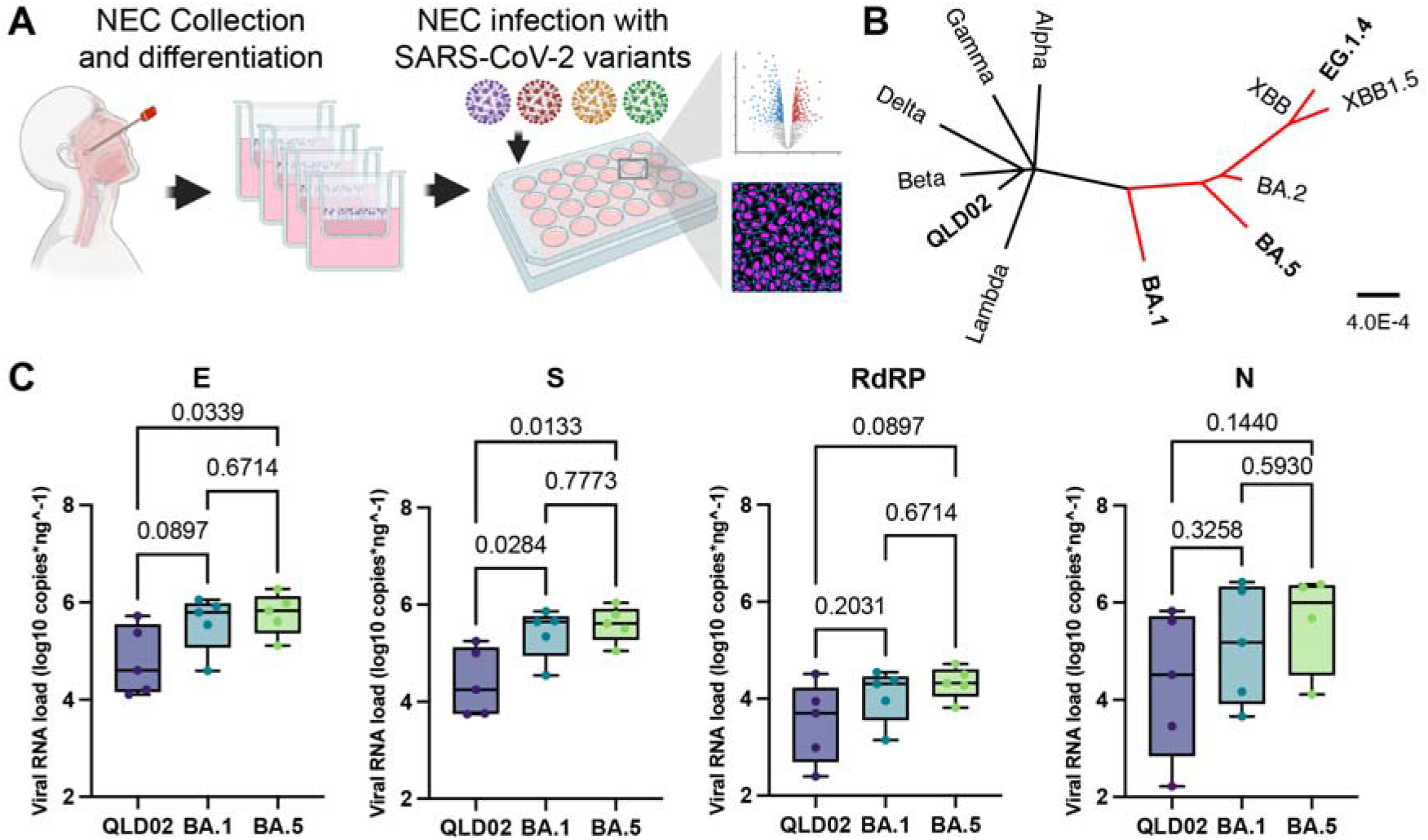
SARS-CoV-2 BA.1 and BA.5 Omicron variants replicate more efficiently in primary human nasal epithelial cells than the ancestral QLD02 isolate. (A) Schematic overview of experimental workflow. Primary human nasal epithelial cells (NECs) were collected from healthy donors and differentiated at air-liquid interface for 28 days to establish pseudostratified ciliated epithelium. Differentiated NECs were subsequently infected and characterised through RNA-seq and immunofluorescence staining. (B) Whole genome phylogenetic inference showing evolutionary relationships among SARS-CoV-2 variants. The Omicron variants are highlighted in red, with isolates examined in this study shown in bold (QLD02/ancestral, BA.1, BA.5, XBB (EG.1.4)). Scale bar indicates nucleotide substitutions per site. (C) RT-qPCR analysis of viral gene expression (E, S, RdRP, and N) in NECs infected with QLD02 SARS-CoV-2 ancestral isolate and Omicron variants BA.1 and BA.5 at 48 hours post-infection. Viral RNA loads are shown as log_10_ copies per ng of total RNA. Each data point corresponds to a NEC culture generated from an independent donor. Error bars indicate median and interquartile range. Statistical significance was determined using the Kruskal-Wallis test.

### Different variants of SARS-CoV-2 elicit distinct transcriptional responses in human primary nasal epithelial cultures

Having established that Omicron variants exhibit enhanced intracellular replication relative to ancestral virus (Fig. 1C), we sought to understand the corresponding impact on the host transcriptional landscape. Total RNA from differentiated NEC cultures at 48 hours post-infection was used for bulk RNA-seq. Analysis of the host response (Fig. S1B, C) revealed distinct transcriptional responses following infection with SARS-CoV-2 variants at 48 hours post infection. Differential expression analysis identified 189, 325, and 4,839 significantly differentially regulated genes (FDR < 0.05) in QLD02-, BA.1-, and BA.5-infected cells relative to mock (Tables S2-S4), respectively (Fig. 2A). Strongly upregulated genes included canonical interferon-stimulated genes (ISGs) such as IFIT1, IFIT2, IFIT3, MX1, MX2, OASL, and IFNB1, alongside pro-inflammatory chemokines CXCL10 and CXCL11 with the magnitude of ISG induction broadly similar across variants. Additionally, a subset of genes encoding ciliary structural and motor proteins were downregulated across infections, with the most extensive and coordinated downregulation signature observed in BA.5-infected cells.

**Figure 2.**
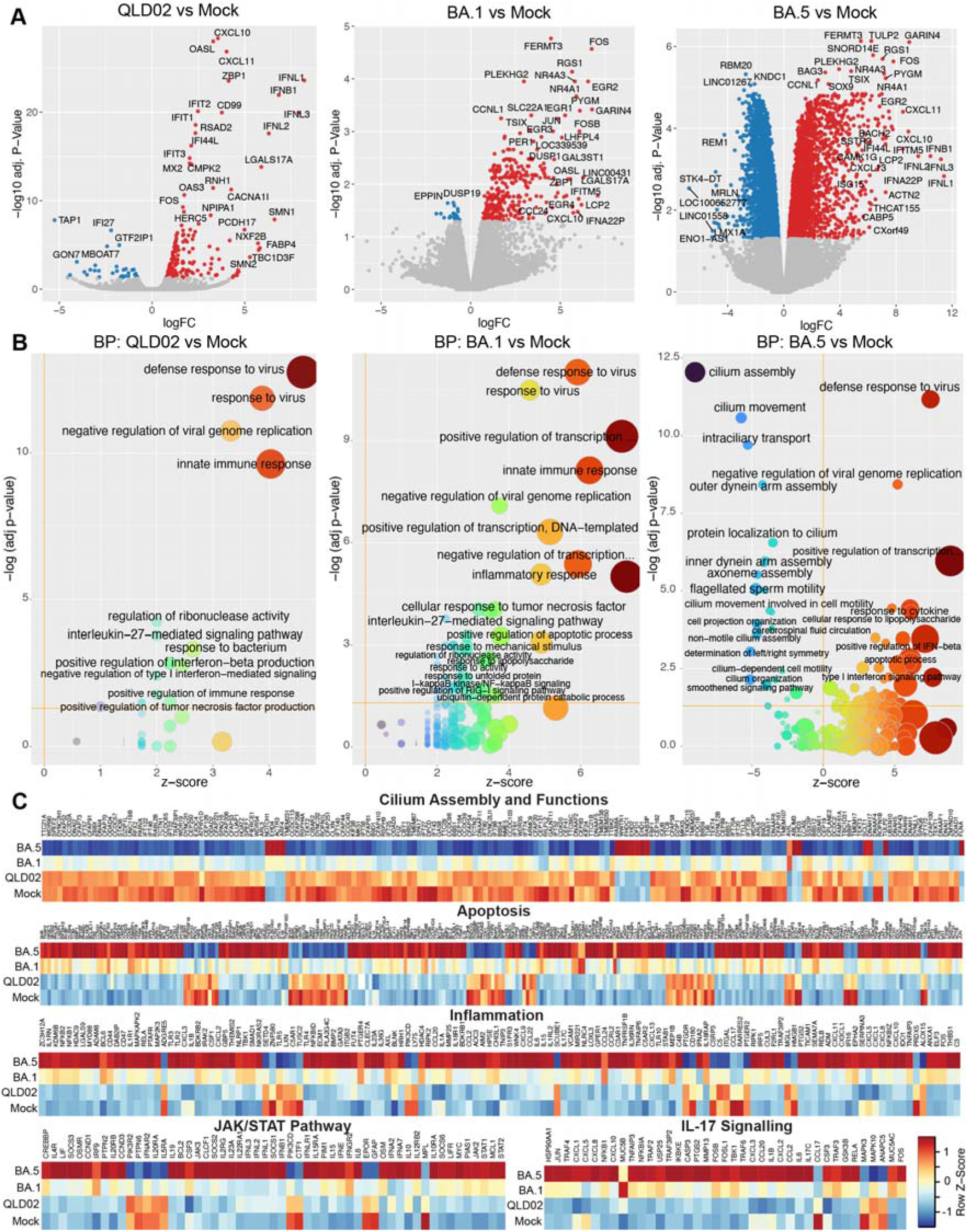
Infection with BA.5, but not BA.1 or the ancestral isolate of SARS-CoV-2 drastically decreases levels of transcripts associated with cilia function and increases apoptotic and inflammatory responses in human NECs. (A) Differentially expressed genes (DEGs) in primary nasal epithelial cells infected with ancestral isolate (QLD02), and Omicron variants (BA.1, and BA.5) of SARS-CoV-2 relative to mock-infected cells. Significantly (FDR-corrected p-value <0.05, log_2_FC>1) upregulated genes are shown in red and downregulated genes are shown in blue. The data are derived from the primary cell cultures obtained from four donors. (B) Gene Ontology (Biological Process) enrichment analysis of DEGs identified in each comparison shown in (A). Z-scores indicate cumulative activation or suppression of biological processes. Size of the bubbles is proportional to the number of DEGs associated with the respective GO term. (C) Heatmaps representing relative expression of the genes associated with cilium assembly and function, apoptosis, inflammation, JAK/STAT signaling, and IL-17 signaling in mock-, QLD02-, BA.1-, and BA.5-infected NECs. The values are the means from 4 individual donors.

Gene Ontology (GO) enrichment analysis highlighted robust activation of innate immune pathways, including “defense response to virus,” “type I interferon signaling,” and “negative regulation of viral genome replication” across all variants (Fig. 2B). BA.5 infection elicited the most distinctive transcriptional response, dominated by a downregulation of cilium-related terms, including “intraciliary transport” and “outer dynein arm assembly”, with the GO term “cilium assembly” (FDR-corrected p-value = 8.96E-13) emerging as the most significantly enriched term. These cilium-related processes were minimally affected in QLD02 and BA.1 infections.

Heatmap visualization of the most significantly affected pathways revealed coordinated repression of cilium assembly and motility genes in both BA.1 and BA.5, with BA.5 exhibiting the greatest reduction (Fig. 2C). Within the cilium assembly and functions gene set, BA.5-infected cells displayed widespread downregulation of structural and motor protein-encoding genes that were relatively preserved in QLD02 and, to a lesser extent, BA.1 infections. This suppression encompassed core structural and motor protein genes of the ODA, including dynein heavy and intermediate chains (DNAH5, DNAH9, DNAI1, DNAI2) (11), intra-flagellar transport genes (IFT88, IFT172), and ciliary assembly factors (CFAP43, CFAP53). AGR3, a ciliated-cell-restricted regulator of calcium-dependent ciliary beat frequency (20), was similarly downregulated (logFC = −1.46, FDR < 0.01), indicating that BA.5 compromises both axonemal force generation and its regulatory control. Both BA.1 and BA.5 also induced more pronounced upregulation of pro-apoptotic factors compared to QLD02 (Fig. 2C) including extrinsic pathway death-receptor ligands (TNFSF10, TNFSF13B) and downstream caspases (CASP1, CASP10), with the strongest induction in BA.5.

Progressive upregulation of inflammation-associated chemokines (CXCL2, CXCL8, CCL20) and cytokines (IL1A, IL1B, IL6), core JAK/STAT pathway components (JAK2, STAT1, STAT2) and IL-17 signalling mediators (NFKB1, CXCL8, CCL20) showed the strongest upregulation in BA.5-infected cells, consistent with enrichment of IL-17-associated transcriptional programmes. Together, these data demonstrate that while all SARS-CoV-2 variants elicit canonical antiviral interferon programs in differentiated nasal epithelia, BA.5 induces a distinctive transcriptomic signature defined by suppression of ciliary gene expression and heightened inflammatory and apoptotic signalling.

### BA.5 variant shows enhanced disruption of ciliary processes, and greater transcription signatures of apoptosis compared to BA.1

To further elucidate the effect of BA.5 infection on cilia disruption we performed a differential gene expression analysis comparing BA.5 and BA.1 infections. This comparison revealed 1,513 significantly differentially expressed genes (FDR < 0.05, |log_2_FC| > 1; Fig. 3A, Table S5). Gene Ontology enrichment analysis of these DEGs (Fig. 3B) revealed key differences in the activation and suppression of pathways associated with apoptosis and ciliary processes.

**Figure 3.**
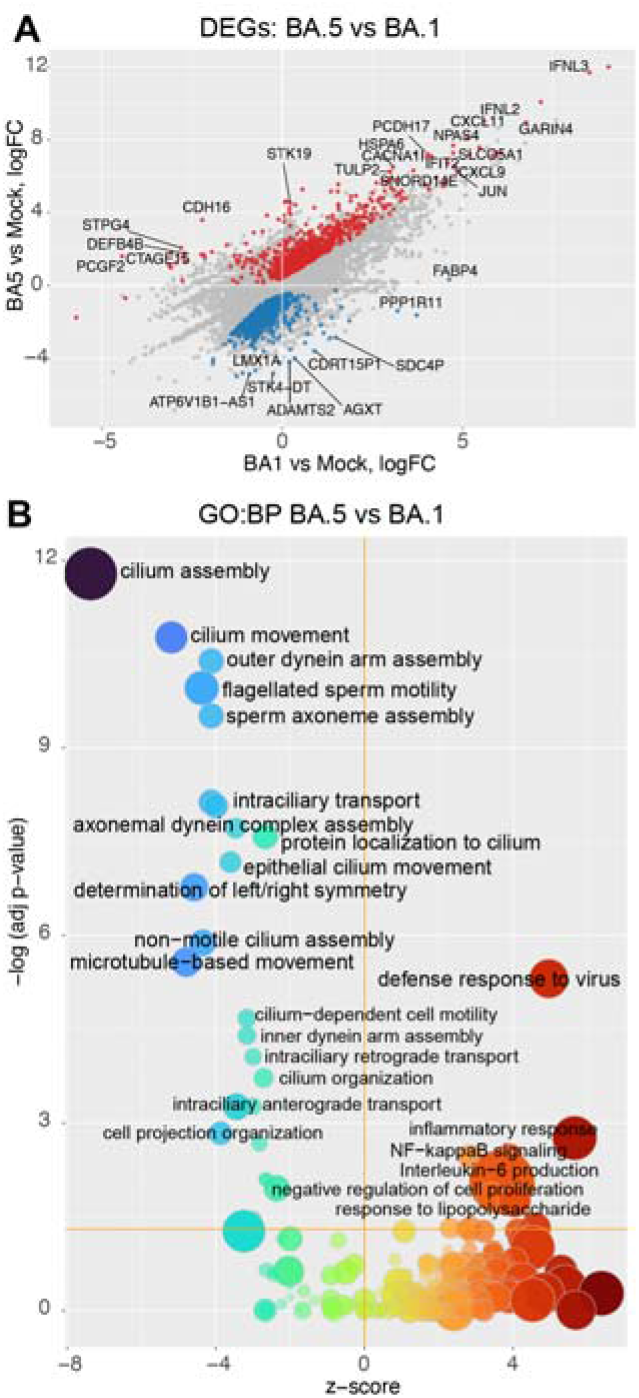
The BA.5 variant elicits stronger antiviral responses and causes significantly more profound disruption to cilia function compared to BA.1. (A) Differentially expressed genes (DEGs) in human primary nasal epithelial cells infected with BA.5 variant of SARS-CoV-2 in comparison to BA.1 infection at 48 hours post-infection. Significantly upregulated genes (log2FC > 1, FDR < 0.05) are shown in red and significantly downregulated genes (log2FC < -1, FDR < 0.05) are indicated in blue. Topmost differentially expressed genes are labelled. (B) Gene Ontology Biological Process (GO:BP) enrichment analysis of the differentially expressed genes identified in BA.5 versus BA.1 comparison shown in (A). Bubble size corresponds to number of DEGs per pathway, colour gradient represents z-score (red = activation, blue = suppression).

First, BA.5 showed a pronounced suppression of cilium-related biological processes, with highly negative z-scores for “cilium assembly” (z-score = -4.2), “cilium movement” (z-score = -3.8), “outer dynein arm assembly” (z-score = -3.5), and “intraciliary transport” (z-score = -3.2). These pathways were minimally affected in the BA.1 versus mock comparison (Fig. 2B), indicating that this ciliary shutdown is largely specific to BA.5.

The BA.5 and BA.1 comparison also revealed a strong positive enrichment (z-scores > 4) for pathways related to “defense response to virus” and “inflammatory response,” which included a coordinated induction of apoptosis-associated genes (Table S5). A closer analysis revealed the upregulation of numerous TNF-superfamily ligands and death-receptor–associated transcripts in BA.5 compared to BA.1, including TNFSF10 (TRAIL; logFC = 1.36) and TNFSF13B (BAFF; logFC = 2.10), along with downstream apoptotic effectors like CASP10, CASP1, and BAK1. Gene Set Enrichment Analysis (GSEA) further confirmed the significant enrichment of apoptosis-related pathways, such as KEGG Apoptosis and Reactome death-receptor signalling pathways (adj. p-values 8×10^−3^ to 3×10^−2^). These signatures suggest a connected biological program in which TNF-family ligands (e.g., TNFSF10) initiate death-receptor signalling upstream of caspase activation. Together these data demonstrate that the divergent pathogenesis of BA.5 compared to BA.1 is defined by a selective suppression of ciliary gene programs and robust activation of TNF-mediated apoptotic signalling.

### XBB has a similar effect to BA.5 on host gene expression in NECs

We next sought to investigate whether Omicron variants that emerged after BA.5 (e.g. XBB) also displayed global downregulation of cilia-related transcriptional processes. Accordingly, NECs from five independent donors were infected with the XBB and BA.5 variants. At 48 hours post-infection, total RNA was isolated from SARS-CoV-2- and mock-infected cells and analysed by RNA-Seq (Fig. 4, Fig. S2), yielding comparable library sizes across conditions (Fig. S2A). XBB replicated at levels comparable to BA.5 (Fig. 4A). Transcriptional profiling of the host gene expression followed by the multidimensional scaling (MDS) analysis demonstrated strong separation of infected groups from mock-infected controls, but no clear separation between BA.5 and XBB (Fig. S2B). Although the donor-stratified MDS analysis revealed subtle separation between BA.5 and XBB infections (Fig. S2C), the overall clustering patterns suggest minor transcriptomic divergence of the cells infected with BA.5 and XBB variants.

**Figure 4.**
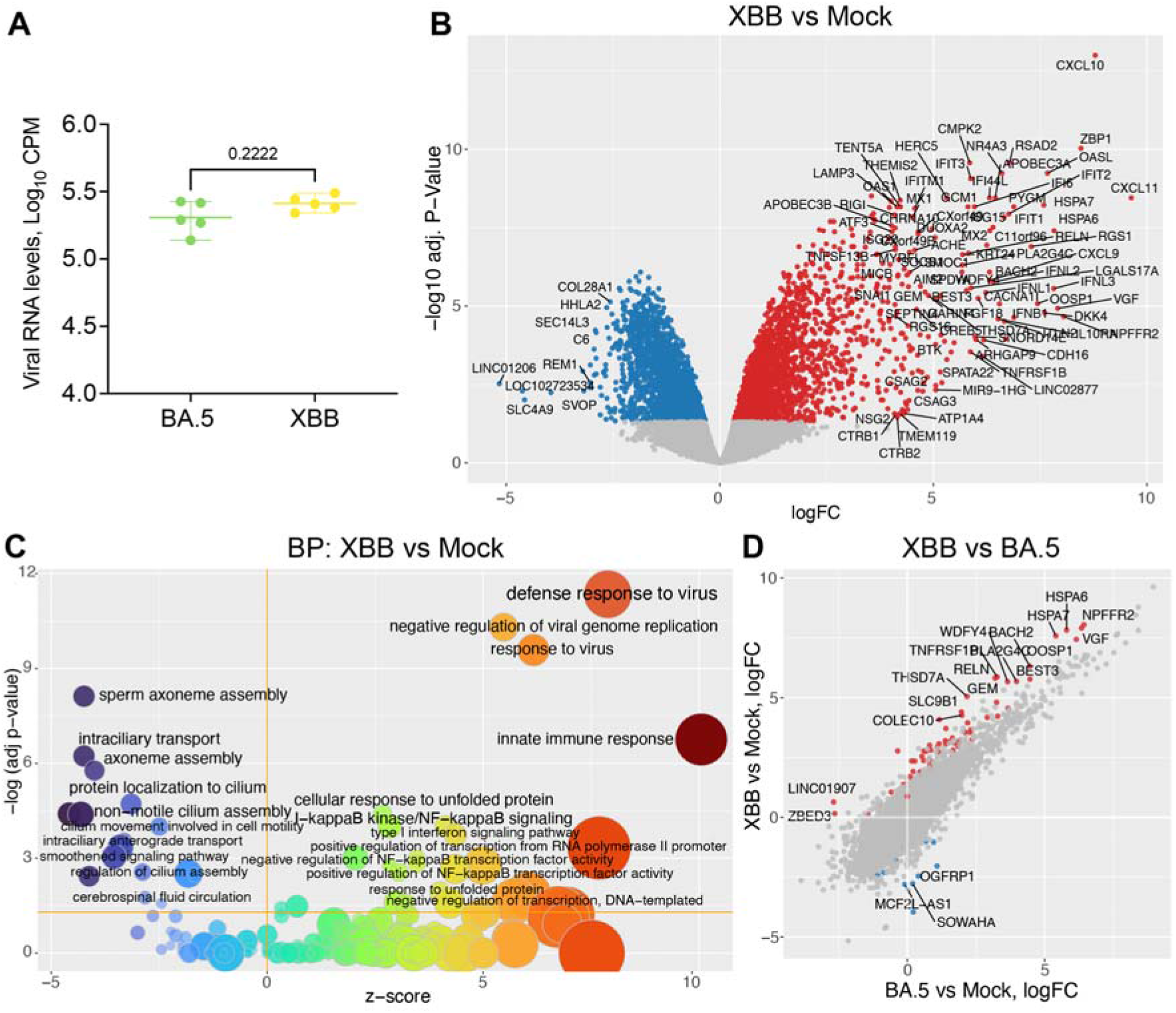
The Omicron XBB (EG.1.4) variant demonstrates comparable replication to BA.5 and results in similar transcriptional profile in primary human nasal epithelial cells. (A) Viral RNA levels in human NECs at 48 hours after infection with XBB and BA.5 SARS-CoV-2 variants. Viral RNA levels were determined by RNA-Seq. Each data point represents an independent cell culture derived from a different donor (n = 5). Bars indicate mean and interquartile range. Statistical significance determined using Mann-Whitney test. (B) Differentially expressed genes (DEGs) in XBB-infected versus mock-infected primary NECs at 48 hours post-infection. (C) Gene Ontology Biological Process (GO:BP) enrichment analysis of XBB infection versus mock-infected control. Bubble size corresponds to number of DEGs per pathway, color gradient represents z-score (red = activation, blue = suppression). (D) Differentially expressed genes (DEGs) in human NECs infected with XBB SARS-CoV-2 variant in comparison to BA.5 infection. In (B) and (D) significantly upregulated genes (log2FC 1, FDR < 0.05) are shown in red and significantly downregulated genes (log2FC < -1, FDR < 0.05) are indicated in blue. Most significantly differentially expressed genes are labelled.

Differential gene expression analysis comparing XBB-infected cultures to mock-infected controls identified a robust antiviral and inflammatory transcriptional response (Fig. 4B), characterized by strong upregulation of canonical ISGs including IFIT1, IFIT2, IFIT3, MX1, MX2, OASL, and RSAD2, alongside elevated expression of chemokines CXCL10, CXCL11, and CCL20. Similar induction of these genes was observed during BA.5 infection (Fig. 2A, Fig. S2D), demonstrating conservation of the antiviral response between the two Omicron variants. Consistent with BA.5 infection, XBB-infected NECs displayed a large number of downregulated genes (BA.5: 3,142 genes; XBB: 4,502 genes; Tables S6-S7). GO enrichment analysis confirmed that downregulated genes were associated with cilia-related processes (Fig. 4C). Key structural and motility-associated transcripts (DNAH5, DNAH9, DNAI1, IFT88, CFAP43; Tables S6-S7) and the calcium-dependent ciliary beat regulator AGR3 were consistently downregulated, reflecting a conserved transcriptomic signature of ciliary motor and regulatory dysfunction in both XBB and BA.5 variants. Direct comparison of XBB and BA.5 infections revealed minimal transcriptional divergence (Fig. 4D) with 125 DEGs meeting significance threshold (Table S8).

XBB therefore elicits a host response highly conserved relative to BA.5, suggesting that late Omicron variants have converged on a similar programme of ciliary dysfunction and antiviral activation.

### BA.1 and BA.5 infections are associated with elevated apoptosis in differentiated NECs without altering overall number of ciliated cells

Our transcriptomic analyses (Figs. 2-4) consistently demonstrated that BA.5 and the later XBB variant, induce upregulation of pro-apoptotic pathways and downregulation of genes essential for ciliary function. To investigate the cellular basis for these transcriptional changes, we used immunofluorescence staining to dissect cell-specific tropism and apoptosis. NECs infected with QLD02, BA.1, or BA.5 were stained for acetylated α-tubulin (cilia, green), cleaved caspase-3 (CC3; apoptotic cells, magenta) and viral nucleocapsid protein (NP; infected cells, orange).

The XY (Fig. 5A) and XZ (Fig. 5B) projections revealed clear differences between variants. In BA.1- and BA.5-infected cultures, there was a greater abundance of both infected (NP^+^) and apoptotic (CC3^+^) cells compared to QLD02 and mock-infected controls. XZ projections confirmed that the cilia (green) form the apical layer of the epithelium and that this is the primary site of viral infection (orange). CC3^+^ staining (magenta) was prominent in the BA.1- and BA.5-infected NECs, with many apoptotic cells in proximity to, but distinct from, NP^+^ cells, suggesting a bystander apoptotic effect.

**Figure 5.**
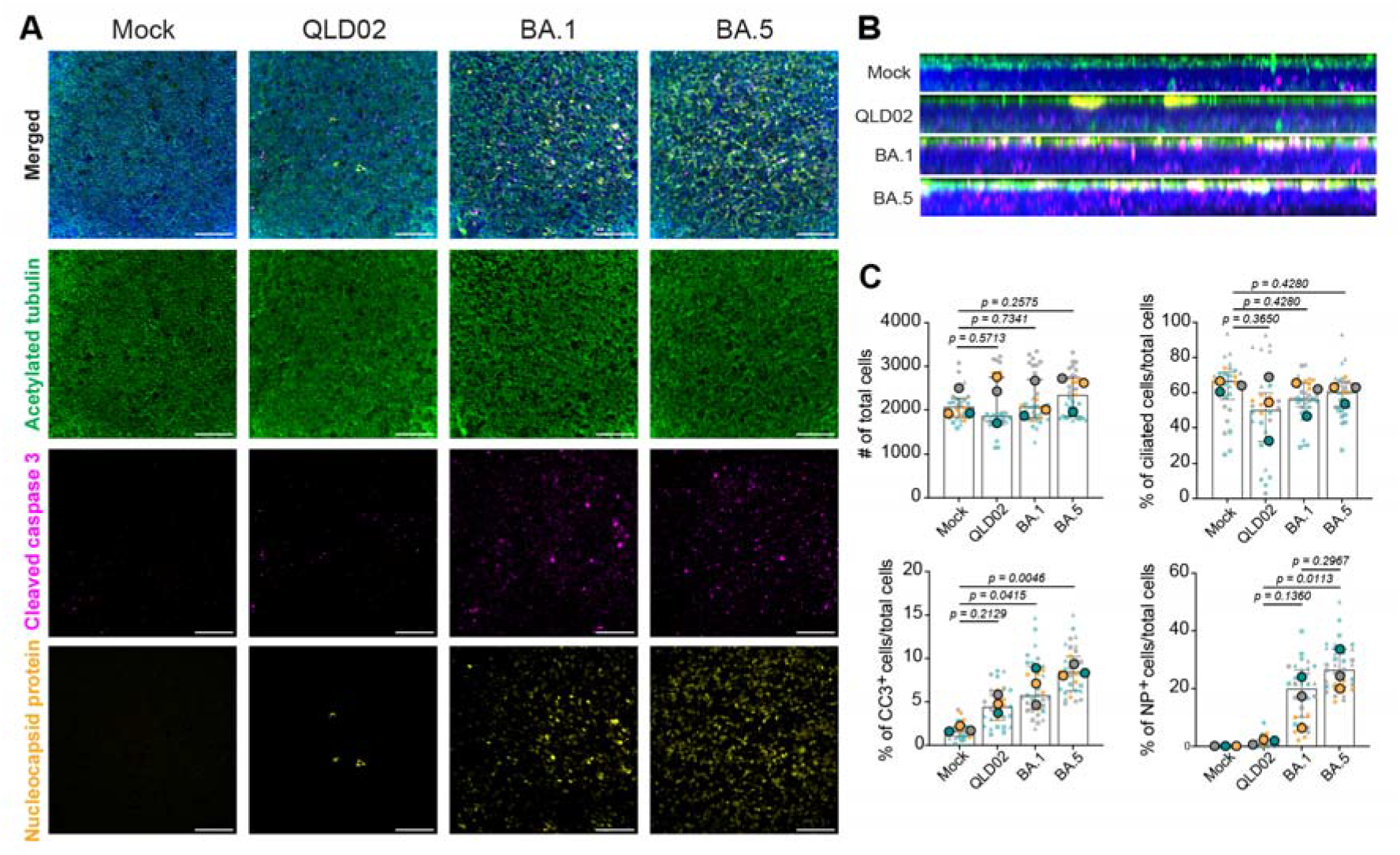
BA.5 infection induces elevated apoptosis and higher viral burden in primary nasal epithelial cells. (A) Representative maximum-intensity projection immunofluorescent microscopy images of differentiated primary NECs infected for 48 h with ancestral QLD02, Omicron BA.1, or Omicron BA.5 variants. Cells were stained for nuclei (DAPI, blue), acetylated α-tubulin (green, cilia), cleaved Caspase-3 (CC3; magenta, apoptotic cells), and nucleocapsid protein (NP; orange, infected cells). Scale bars = 100 µm. (B) Maximum-intensity projections of (A) in the XZ plane. (C) Quantification of total number of DAPI^+^ cells, acetylated α-tubulin^+^ cells (% ciliated cells), CC3^+^ cells (% apoptotic cells) and NP^+^ cells (% infected cells) across independent donors and replicate cultures. Each small point represents a single field of view (n = 5 per replicate). Large points represent the means of 1-3 replicates per each donor, with 3 independent donors indicated by different coloured data points. Statistical significance was determined using the Kruskal-Wallis test; pairwise comparisons were made with Uncorrected Dunn’s test. Error bars represent median with interquartile range.

Quantification (Fig. 5C) confirmed no differences in total cells across conditions. BA.5 showed a significantly higher percentage of infected (NP^+^) cells (median, 23.99%) compared to QLD02 (median, 1.83%), representing a 13.11-fold increase (p = 0.0113). The 9.33-fold increase for BA.1 (median, 17.07%), relative to QLD02 was not statistically significant (p = 0.1360).

Increases in the percentage of infected cells were accompanied by similarly increased percentages of apoptotic (CC3^+^) cells across BA.1 and BA.5 infection. BA.1 infection (median, 7.044%) showed a 4.20-fold increase in CC3+ cells (p = 0.0415), and BA.5 infection (median, 8.254%) a 4.92-fold increase (p = 0.0046), relative to mock (median, 1.677%). However, despite the profound downregulation of ciliary genes seen in the transcriptomic data, there was no variant-dependent difference in the total number of ciliated cells at 48 hours post-infection (Fig. 5C). This suggested that the BA.5-associated phenotype may reflect early functional impairment of structurally retained cilia, rather than overt cilia loss.

### BA.5 and BA.1 Omicron variants are associated with increased infection and apoptosis of ciliated cells, relative to the ancestral SARS-CoV-2 virus

To further dissect the interactions between infection, apoptosis, and ciliation, we examined colocalization by immunofluorescence (Fig. 6). Representative images (Fig. 6A) revealed strong overlap between infected (NP^+^) and apoptotic (CC3^+^) cells, in BA.1 and BA.5, with increased double-positive infected (NP^+^) / ciliated (α-tubulin^+^) cells, indicating enhanced tropism for ciliated cells.

**Figure 6.**
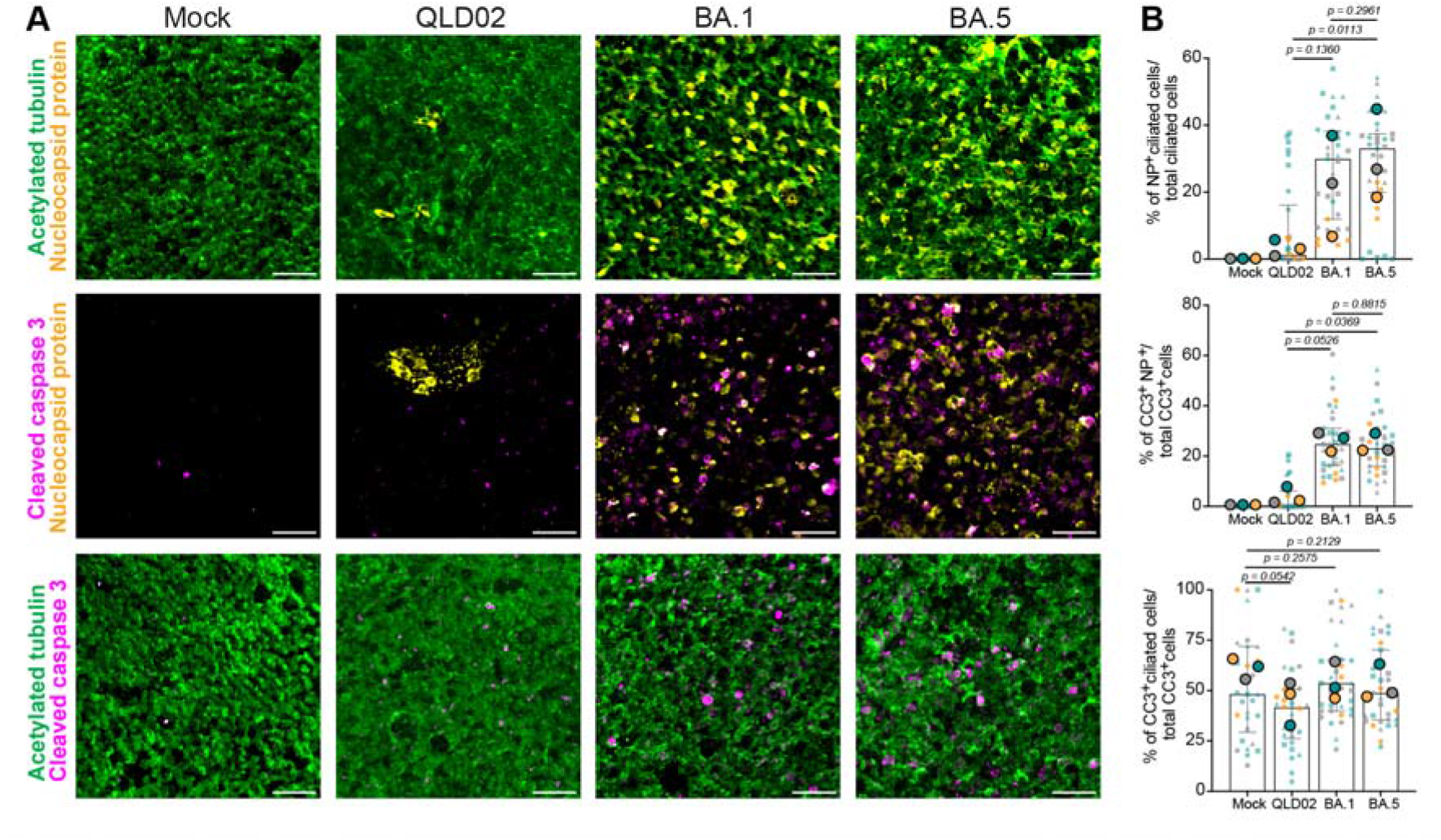
BA.5 and BA.1 infections of NECs are associated with increased apoptosis of infected cells and preferential ciliated cell tropism compared to QLD02. (A) Representative maximum-intensity projection immunofluorescent microscopy images of differentiated primary NECs infected for 48 h with ancestral QLD02, Omicron BA.1, or Omicron BA.5 variants. Cells were stained for nuclei (DAPI, blue), acetylated α-tubulin (green, cilia), cleaved Caspase-3 (CC3; magenta, apoptotic cells), and nucleocapsid protein (NP; orange, infected cells). Scale bars = 50 µm. (B) Quantification of the percentage of infected ciliated (NP^+^ acetylated α-tubulin^+^) cells (top) over total ciliated cells, all apoptotic infected (CC3^+^ NP^+^) cells over total CC3^+^ cells (middle), and percentage of apoptotic ciliated (CC3^+^ acetylated α-tubulin^+^) cells over total CC3^+^ cells (bottom). Each small point represents a single field of view (n = 5 per replicate). Large points represent the means of 1-3 replicates per each donor, with 3 independent donors indicated by different coloured data points. Statistical significance was determined using the Kruskal-Wallis test; pairwise comparisons were made with Uncorrected Dunn’s test. Error bars represent median with interquartile range.

Quantitative analysis first confirmed that for Omicron variants (BA.1 and BA.5), there was an enhanced tropism for ciliated cells compared to the ancestral virus (QLD02). BA.5-infected NECs showed a significantly higher percentage of infected ciliated cells (NP^+^ / acetylated α-tubulin^+^, median, 25.73%) compared to QLD02 (median, 2.87%), representing an 8.97-fold increase (Fig. 6B; p = 0.0113). BA.1 also showed a 7.55-fold increase in double-positive infected ciliated cells (median, 21.66%), relative to the QLD02 strain; however, this difference did not reach statistical significance (p = 0.1360).

Next, we directly linked the infection status (NP^+^) of a cell to its apoptotic status (CC3^+^). In Omicron-infected cultures, the fraction of double-positive apoptotic and infected (CC3^+^ and NP^+^) cells was 3.77-fold higher for BA.5 (median, 28.34%) compared to QLD02 (median, 7.52%, Fig. 6B; p = 0.0369). BA.1 showed a 3.22-fold increase (median, 24.22%) over the QLD02 virus; however, this comparison did not reach statistical significance (p = 0.0526). There was no significant difference observed between the BA.1 and BA.5 variants in this regard (p = 0.8815). Despite increases in apoptosis of infected cells and increased tropism for ciliated cells, there were no statistically significant changes in the fraction of apoptotic ciliated (CC3^+^ α-tubulin^+^) cells, across all variants (Fig. 6B).

Omicron variants (BA.1 and BA.5) therefore exhibit enhanced tropism for ciliated cells (Fig. 6B, top panel) that is associated with an increase in apoptosis of infected cells; but not ciliated cells specifically, suggesting a bystander apoptotic effect distinct from the ancestral virus.

### BA.5 infection reduces DNAH5 outer dynein arm protein in ciliated nasal epithelial cells

A key limitation of studying SARS-CoV-2 pathogenesis under BSL-3/PC3 containment is the difficulty to perform live-cell ciliary beat frequency or mucociliary transport assays on infected cultures. To obtain a functional readout of ciliary motor function, we quantified ciliary DNAH5 by immunofluorescence. DNAH5 is a core heavy-chain component of the outer dynein arm, the axonemal motor complex that powers ciliary beating (10). Reduced DNAH5 at the ciliary axoneme is therefore expected to reflect impaired ciliary motor capacity even when cilia remain morphologically intact (11). Differentiated NECs at 48 hours post-infection were stained for DNAH5, acetylated α-tubulin (total cilia area), and DAPI (nuclei) (Fig. 7A). DNAH5 fluorescence intensity normalised to total cilia area (IntDen/Cilia Area) was comparable across mock, QLD02, and BA.1 (Fig. 7B). In contrast, BA.5-infected NECs showed a significant reduction in DNAH5 protein density relative to both mock (p = 0.0092) and BA.1 (p = 0.0415). This reduction occurred without a corresponding decrease in total ciliated cell number (Fig. 5C), indicating that the BA.5-associated loss of DNAH5 reflects a selective depletion of outer dynein arm complexes from morphologically intact cilia. Donor-matched normalised RNA-seq data from two independent experiments (Figs 2,4) showed reduced DNAH5 transcript abundance in both BA.5- and XBB-infected NECs relative to QLD02 (p<0.05, Kruskal-Wallis test), with no difference between BA.5 and XBB (Fig. 7C).These data demonstrate that the BA.5 transcriptomic signature of ciliary gene downregulation translates to measurable loss of a key motor protein required for ciliary beat generation, indicative of impaired mucociliary clearance in BA.5-infected nasal epithelia.

**Figure 7.**
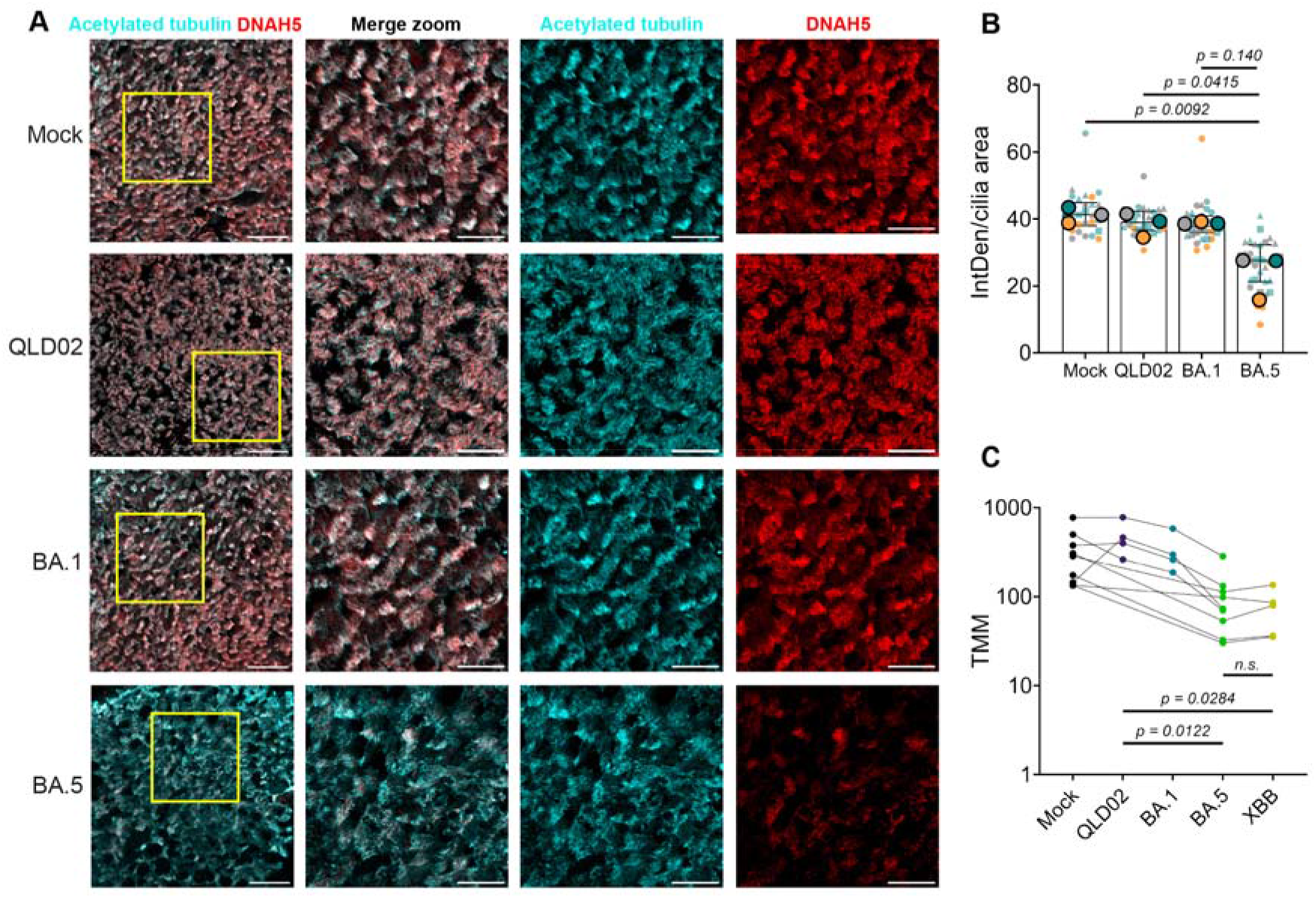
BA.5 infection reduces DNAH5 outer dynein arm protein expression in ciliated nasal epithelial cells. (A) Representative maximum-intensity projection immunofluorescent microscopy images of the apical cilia region of differentiated primary NECs infected for 48 h with mock (PBS), ancestral QLD02, Omicron BA.1, or Omicron BA.5 variants. Cells were stained for acetylated α-tubulin (cyan, cilia) and DNAH5 (red, outer dynein arm heavy chain). Scale bars = 50 µm, scale bars of zoomed inset = 25 µm. (B) Quantification of DNAH5 fluorescence intensity normalised to total cilia area (IntDen/Cilia Area) across infection conditions. Each small point represents a single field of view (n = 5 per replicate). Large points represent the means of 1–3 replicates per donor, with 3 independent donors indicated by different coloured data points. Statistical significance was determined using the Kruskal-Wallis test with uncorrected Dunn’s post-hoc test. Error bars represent median with interquartile range. (C) TMM (Trimmed Mean of M-values) -normalised expression of DNAH5 with lines connecting matched donors across the two independent RNA-seq experiments. Statistical significance was determined by Kruskal-Wallis test with Dunn’s multiple comparisons test.

## Discussion

SARS-CoV-2 Omicron has evolved increased replication efficiency in the upper respiratory tract compared to the ancestral virus and Delta variant, with evidence suggesting enhanced binding to motile cilia in human nasal epithelial cells (13). However, despite the emergence of multiple Omicron variants (e.g. BA.1, BA.5, XBB), no studies to date have systematically assessed how these variants differ in their interaction with and impact on ciliated cells, leaving a gap in our understanding of disease pathogenesis associated with more recent Omicron variants.

Here, we show that whilst ancestral SARS-CoV-2 (QLD02), BA.1 and BA.5 induced numerous transcriptional changes in primary human NECs, the most striking difference between Omicron variants was a pronounced downregulation in pathways for cilia function such as ‘cilium assembly’ and ‘cilium movement’ in BA.5-infected cells, relative to BA.1 infected cells. This was accompanied by increased expression of proinflammatory genes (e.g. ‘defence response to virus’) and increased activation of apoptosis pathways (e.g. ‘positive regulation of apoptotic processes’) relative to BA.1 infected cells. It would therefore be tempting to speculate that this transcriptional phenotype in BA.5 is driven by increased death of ciliated cells and subsequent loss of cilia. However, immunofluorescence staining revealed that this was not the case. Whilst BA.5 did cause increased apoptosis (likely accounting for the increased inflammation seen with this strain by transcriptomics) this was not more pronounced in ciliated cells in any variant tested (including the ancestral virus). Similarly, there was no difference in total ciliated cell populations between the ancestral virus and Omicron variants at 48 hours post-infection. These data are broadly consistent with prior studies suggesting that both the ancestral virus (7) and Omicron variants (18) do not alter tubulin staining in the first 48 hours of infection. Our transcriptomic results are also corroborated by functional evidence that SARS-CoV-2 can decrease cilia beat frequency and coordination in the absence of widespread cilia loss (7).

Importantly, ciliary DNAH5 staining further clarifies the BA.5 phenotype. Although acetylated α-tubulin staining showed that the overall ciliated layer remained morphologically intact at 48 hours post-infection (Fig. 5C), BA.5 significantly reduced DNAH5 signal when normalised to cilia area (Fig. 7B). Because DNAH5 is a core outer dynein arm motor protein required for effective ciliary beating (11), this finding suggests that BA.5 impairs the motility machinery of retained cilia rather than causing loss of the ciliary compartment. A consequence of this phenotype could be reduced mucociliary clearance, a major innate defence mechanism of the upper airway in which mucus and coordinated ciliary beating act together to trap and remove inhaled particles and pathogens (21). Impairment of this clearance could favour viral persistence at the epithelial surface, particularly given that SARS-CoV-2 preferentially targets cilia and may use ciliated cells to facilitate access to the underlying epithelium (7, 13). In this context, impaired ciliary motility may help explain why BA.5 produced higher viral burden than BA.1 and QLD02 in our NEC model (Fig. 1C). When combined with the increased mucus production reported during SARS-CoV-2 infection (22), we hypothesize that reduced cilia motility would impair mucociliary transport, resulting in the accumulation and thickening of airway secretions. This stagnant mucus layer can retain viral particles and proinflammatory cytokines at the epithelial surface, prolonging viral exposure, exacerbating local inflammation, and facilitating continued infection and spread. Together, these data support a model in which BA.5 induces early ciliary motor dysfunction before overt deciliation, with likely consequences for mucosal barrier function and viral replication.

We further suggest that the increased apoptosis observed in BA.5-infected cells reflects predominantly the death of other non-ciliated cells in the differentiated epithelium (e.g. goblet and/or basal cells). Indeed, these data are consistent with previous studies showing a “bystander apoptotic effect” where necroptosis is predominately activated in SARS-CoV-2 infected epithelial cells whilst neighbouring cells undergo apoptosis (23).

The question therefore remains as to why the phenotype of cilia dysfunction, increased apoptosis of non-ciliated cells and increased inflammation is more pronounced in BA.5 and XBB, compared to earlier SARS-CoV-2 variants. Importantly, these trends are not the result of differences in viral replication efficiency, as BA.1, BA.5 and XBB had similar viral reads at 48 hours post-infection. Instead, it is possible that these data reflect the increased fusogenicity and syncytia formation of BA.5 compared to earlier Omicron variants (24, 25). Syncytial formation and the resultant altered cytoskeletal dynamics have been shown to interfere with the coordinated orientation and beating of cilia, leading to reduced mucociliary clearance even when cilia remain morphologically intact (7). Syncytial formation in ciliated cells may also account for the apoptosis of uninfected bystander cells (and subsequent inflammation). Direct comparison of syncytial formation in primary human NECs after BA.1, BA.5 and XBB infection thus remains an important area for future study.

Whilst these data were generated in an *in vitro* system, it is important to consider the *in vivo* implications of our observations. When ciliary motility is reduced or discordant *in vivo*, mucus transport is slowed, potentially enabling pathogens such as SARS-CoV-2 to penetrate beyond the conducting airways (7, 26). However, determining whether BA.5 or even XBB is more likely to progress to lower respiratory tract infections *in vivo* is difficult to assess, as site specific evaluation of viral replication is confounded by strain dependent differences in tropism for cells of the upper and lower respiratory tract (15, 18). Accordingly, whilst our transcriptomic and protein-level data are consistent with impaired ciliary motor function, it remains to be determined whether this translates to measurable deficits in mucociliary clearance during clinical BA.5 infection.

The present study was subject to several limitations. Firstly, due to the laborious nature of working with primary human NECs only a limited number of SARS-CoV-2 variants were tested, and it would be informative to observe if similar transcriptional patterns were observed with more recent Omicron variants (e.g. JN.1). Finally, future studies would benefit from performing cilia function and motility assays to better understand the impact of the observed phenotype associated with BA.5/XBB infections, although we acknowledge the difficulties of performing such experiments under biosafety level 3 containment conditions.

Nevertheless, the present study provides the first insight into the interaction of different Omicron variants with cilia in the nasal epithelium.

## Methods

### Human Ethics, Nasal Epithelial Swab Collection and Processing

Primary nasal epithelial cells (NECs) were collected from healthy adult volunteers as described previously (27). The study was approved by the University of Queensland Human Research Ethics Committee (2017000520 and 2022002162).

Primary NECs were established as previously described (27) and cryopreserved at passage 1 or 2 in freezing medium consisting of fetal bovine serum (FBS) with 10% dimethyl sulfoxide (DMSO). Cells were expanded and maintained in PneumaCult™-Ex Plus medium (STEMCELL Technologies, Canada) and seeded at a density of 4–5

× 10□ cells per transwell insert (6.5-mm polyester membranes, 0.4-µm pore size; Corning Costar, USA). Cultures were maintained in PneumaCult™-Ex Plus until confluent, after which they were “air-lifted” by removing apical medium and replacing the basolateral medium with PneumaCult™-ALI differentiation medium (STEMCELL Technologies). Basolateral medium was refreshed three times per week. Cells were differentiated at the air-liquid interface (ALI) for at least three weeks, until the appearance of ciliated and mucus-producing cells and attainment of a transepithelial electrical resistance greater than 500 Ωcm^2^. Fully differentiated NEC cultures were used for infection and downstream analyses (Table 1).

**Table 1.**
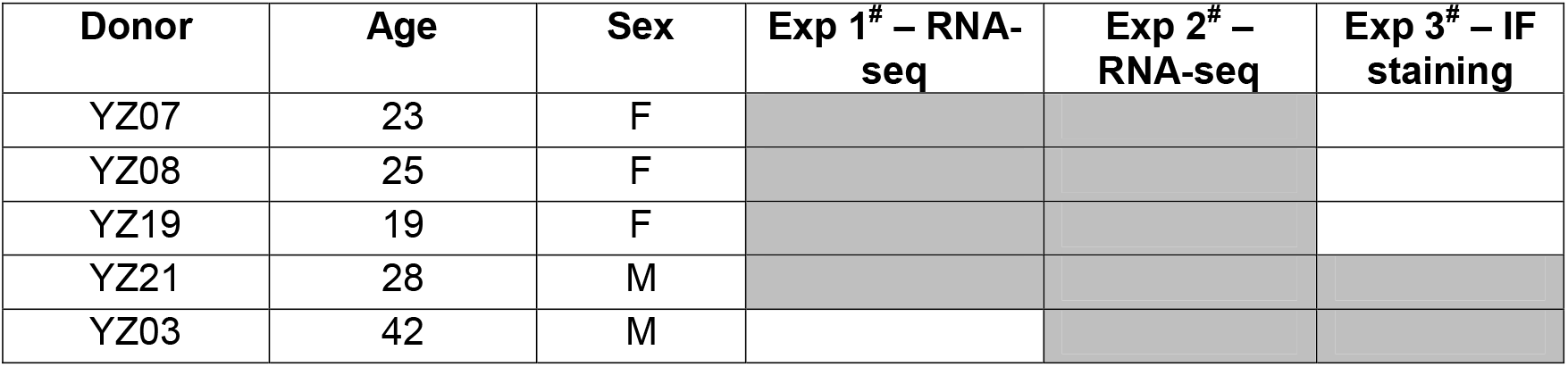
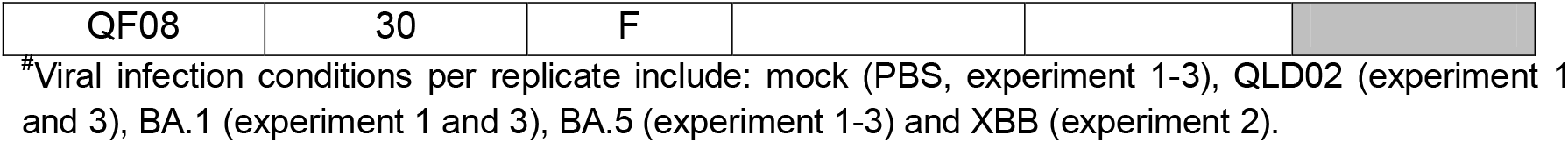
Donor information for primary nasal epithelial cells across three experiments, grey squares indicate donors used in each experiment.

### Cell Lines

VeroE6 stably expressing human TMPRSS2 (VE6-hTMPRSS2 Clone #1), as previously generated and described in (28) were cultured at 37°C and 5% CO_2_ in Dulbecco modified Eagle medium (DMEM; Gibco, Thermo Fisher, USA) supplemented with 10% fetal calf serum (FCS; Bovogen; USA), 1% GlutaMAX (Cat: 35050079, Thermo Fisher, USA), and 100×U/mL penicillin-streptomycin (Gibco, Thermo Fisher, USA), with addition of 30×µg/mL of puromycin dihydrochloride (A1113803, Thermo Fisher, USA).

### SARS-CoV-2 Viruses

Four low-passage SARS-CoV-2 viruses were recovered from nasopharyngeal aspirates of infected individuals provided by the Queensland Health Forensic and Scientific Services, Queensland Department of Health, and the Queensland Institute of Medical Research. These isolates included: (I) an early ancestral (Pango v.4.2 lineage A) Australian isolate hCoV-19/Australia/QLD02/2020 (QLD02) sampled on 30/01/2020 (GISAID Accession ID: EPI_ISL_407896) (28). (II) the Omicron BA.1, named hCoV-19/Australia/QLD2639C/2021 collected on 14/12/2021 (GISAID accession ID: EPI_ISL_8186598) (29); (III) the Omicron BA.5 sub-lineage BE.1 named hCoV-19/Australia/QLD-QIMR03/2022 (GISAID accession ID: EPI_ISL_15671874.2) collected 19/07/2022 (30) and (IV) an XBB.1.9 lineage named hCoV-19/Australia/QLD-UQ01/2023 (GISAID accession ID: EPI_ISL_17784860) collected 11/4/2023 (Unaliased XBB.1.9.2.1.4; Pango lineage EG.1.4) (30). All working virus stocks were propagated and titred on VeroE6 stably expressing human TMPRSS2 (VE6-hTMPRSS2) cells using an immuno-plaque assay (31). All studies with SARS-CoV-2 were performed under physical containment 3 (PC3) conditions and were approved by the University of Queensland Biosafety Committee (IBC/522B/SCMB/2022).

### Viral infection

Differentiated NECs were mock-infected with PBS or infected with QLD02, BA.1, BA.5, or XBB at 1.5 × 10^5^ focus forming units. Specifically, 100 μL of virus or PBS was placed on the epithelial surface in the apical compartment and incubated for 1 hour at 37 °C. Following incubation, excess virus was removed from the transwell, and cells were incubated at 37 °C with 5% CO_2_. Every 24 hours, the basolateral media were refreshed with 1 mL of new ALI media. At predetermined time points, post-infection 100 μL of PBS was added to the apical compartment, and cells were incubated at 37 °C with 5% CO_2_ for 10 minutes. The apical supernatant was subsequently collected and stored at −80 °C.

### Immunofluorescence staining and image quantification

Infected differentiated NECs grown on transwell membranes were fixed with 4% paraformaldehyde (Electron Microscopy Sciences) in PBS for 30 minutes at room temperature and washed three times with PBS. Membranes were then excised from transwell inserts with a scalpel, followed by permeabilization with 0.1% Triton X-100 (Sigma-Aldrich, USA) in PBS for 15 minutes. Samples were then blocked in 1% BSA (Sigma-Aldrich, USA) in PBS for 45 minutes at room temperature. Samples were stained with primary antibodies diluted in 1% BSA in PBS overnight at 4°C. The following primary antibodies were used: mouse anti-acetylated α-tubulin (Cat#66200-1-Ig, Proteintech, USA) as a cilia marker, rabbit anti-cleaved Caspase-3 (Asp175, Cat#9661, Cell Signaling Technology), rabbit anti-DNAH5 (Cat#HPA037470, Sigma-Aldrich), and human anti-SARS-CoV-2 nucleocapsid nanobody E2 (32).

Stained samples were washed three times in 0.5% BSA in PBS, then incubated with secondary antibodies diluted in 1% BSA in PBS for 2.5 hours at room temperature in the dark. Secondary antibodies included: Alexa Fluor 488 donkey anti-mouse IgG (Cat#A32766, Invitrogen, USA), Alexa Fluor 555 donkey anti-rabbit IgG (Cat#A31572, Invitrogen, USA), and Alexa Fluor 647 donkey anti-human IgG (Cat#A21445, Invitrogen, USA). Nuclei were counterstained with DAPI (Invitrogen) for 5 minutes, followed by one wash in PBS. Samples were then briefly rinsed in Milli-Q water, and mounted between two glass coverslips using ProLong Gold Antifade Mountant (Thermo Fisher Scientific, USA).

Z-stack images (∼22 *µ*m, 0.5 *µ*m step size) were acquired on a SpinSR10 spinning disk confocal (Olympus, Japan) using a 30× silicone immersion objective. For DNAH5 staining, z-stack images (∼14 *µ*m, 0.25 *µ*m step size) were acquired using a 60× oil objective. All images were acquired using identical acquisition settings within each objective (30× silicon or 60× oil) and all image quantification was performed using identical analysis parameters within each dataset. All representative images are maximum-intensity image projections generated from Z-stacks. All microscopy imaging was performed at the Queensland Brain Institute, University of Queensland. Cell quantification was performed by acquiring five random fields of view per sample at 2300 × 2300 px resolution, with 1-3 technical replicate samples per group (specified in corresponding figure legends). Image quantification was performed in Fiji/ImageJ (v1.54p) using custom-written scripts to identify and count cells automatically.

Total number of cells were quantified by generating maximum-intensity projections of Z-stacks, applying a Gaussian blur (σ = 2) and automated thresholding on the DAPI channel. Images were converted to a binary mask and DAPI^+^ nuclei detected using Find Maxima (prominence = 20). Point detections were recorded as ROIs and counted automatically using Analyze Particles (<10 µm). Total number of nucleocapsid protein (NP^+^), cleaved Caspase-3 and ciliated (acetylated α-tubulin) cells were quantified using the same workflow as above but applied to their respective fluorescence channels to create a binary mask. The Fill Hole function, followed by Watershed separation was applied to the binary mask, and positive cells were automatically counted using Analyze Particles (< 10 *µ*m).

For quantification of double-positive cells, images were processed as above in the relevant two fluorescence channels. The resulting two binary masks were then combined using ImageJ’s Image Calculator (AND) function to generate an intersection mask representing double-positive cells. Watershed separation was applied to the intersection mask, and double-positive cells were automatically counted using Analyze Particles (<10 *µ*m).

For quantification of ciliary DNAH5 intensity, z-stacks for each field of view were cropped to ∼5 µm to restrict analysis to the apical region containing cilia only, excluding the underlying cell body and nucleus. Maximum intensity projections were then generated from the cropped z-stacks. A binary cilia mask was then generated by thresholding the acetylated tubulin channel. DNAH5 fluorescence intensity within cilia was quantified by applying the binary cilia mask to the DNAH5 channel and measuring DNAH5 mean intensity values within the masked ciliary regions.

### RNA sequencing and analysis

Total RNA was isolated from individual NECs using TRI Reagent (Sigma-Aldrich, USA) as done previously (33). RNA quality and integrity were assessed on an Agilent TapeStation 4200 (Agilent Technologies, USA), and only samples with an RNA integrity number (RIN) > 8 were used for downstream library preparation using the TruSeq RNA Library Preparation Kit v2, Set A (Illumina, USA) according to the manufacturer’s protocol. Indexed cDNA libraries were pooled and sequenced on an Illumina NextSeq 500 instrument using the NextSeq 500/550 High Output Kit v2.5 (75 cycles, Illumina, USA). Image acquisition, processing, and fastq demultiplexing were performed using in-instrument software as described previously (34).

Processing of the raw reads was performed as described previously (35). Raw sequencing reads were assessed for quality using FastQC (v.0.72) and adapter and quality trimmed using Trimmomatic (v.0.36.6) using the parameters: *ILLUMINACLIP:TruSeq3-SE:2:30:10, LEADING:32, TRAILING:32, SLIDINGWINDOW:4:20*, and *MINLEN:16*. High-quality reads were aligned to the human reference genome assembly (GRCh38/hg38) using HISAT2 (v.2.2.1) with default parameters. Gene-level counts were obtained using featureCounts (v2.0.1) with the following options: *counting mode* = “Union,” *strand* = “Unstranded,” *feature type* = “exon,” and *ID attribute* = “gene_id”.

Downstream analysis and visualization of RNA-seq data were conducted in RStudio (v2025.09.2). Differential gene expression analysis was conducted in edgeR (v.4.2). Low abundance reads (<1 count per million) were filtered before normalisation. Library size and composition bias were corrected using the trimmed mean of M-values (TMM) normalisation method. Normalized counts were analysed by multidimensional scaling analysis under default settings and used to build quasi-likelihood negative binomial generalized log-linear model. Contrasts were tested using the quasi-likelihood ratio test or likelihood ratio test with correction for the false discovery rate (FDR < 0.05).

Significantly differentially expressed genes (DEGs) were visualized as volcano plots using ggplot2 (v3.3.2). Gene Ontology (GO) term enrichment was performed using Database for Annotation, Visualization and Integrated Discovery (DAVID, v6.8) as previously described (35). Enrichment scores were integrated with expression values, and z-scores were computed and visualised using GOplot (v.1.0.2) and ggplot2 (v.3.3.2). Heat maps of selected GO Broad Processes (BP) terms were generated using heatmap.2 function of gplots (v3.1.2) based on scaled expression values (row z-scores) (36).

### SARS-CoV-2 Gene Expression Analysis

For quantification of viral genomic RNA abundance, adapter- and quality-trimmed RNA-seq reads as above were aligned to isolate-specific SARS-CoV-2 reference genomes using Bowtie2 (v2.0.5) with default mapping parameters. Read counts were obtained directly from BAM files using samtools v1.18 and normalized as counts per million.

### RNA extraction and quantitative reverse transcription PCR (qRT-PCR)

Total RNA was extracted as described above and used for reverse transcription quantitative PCR (RT-qPCR). For qPCR, 1 μg of isolated RNA was used to generate first-strand cDNA synthesis using the qScript cDNA SuperMix (Quantabio, USA) following the manufacturer’s instructions. Quantitative PCR was conducted on a QuantStudio 6 Flex Real-Time PCR System (Applied Biosystems, USA) using gene-specific primers targeting the SARS-CoV-2 E, S, RdRP, and N genes (Table S1). Each reaction was performed in technical triplicate for each biological replicate. Viral RNA copy number was quantified by comparison to a standard curve generated from serial tenfold dilutions (10^2^ - 10^8^ copies per reaction) of a PCR-amplified and purified SARS-CoV-2 genomic fragment. Copy numbers were normalized to input RNA concentration and expressed as log_10_ copies ng^−1^ total RNA. No-template controls were included in each run.

### Statistical analysis

The number of biological replicates, the sample sizes and statistical methods are indicated in the figure legends. Statistical analysis was performed using GraphPad Prism 10 software. Specific tests included the non-parametric Kruskal-Wallis test with Uncorrected Dunn’s post-hoc test for multiple group comparisons and the Mann-Whitney test for two-group comparisons. RNA-seq analysis was performed using RStudio (v2025.09.2) as detailed in the ‘RNA sequencing and analysis’ section. Statistical significance was defined as p < 0.05.

### Data and materials availability

All data needed to evaluate the conclusions in the paper are present in the paper and/or the Supplementary Materials. RNA sequencing data generated for this study have been deposited in the NCBI Sequence Read Archive (SRA) under accession number PRJNA1347416.

## Author contributions

Conceptualization, A.A.K., K.R.S., R.H.P., and A.S.; Investigation, R.H.P., H.M., G.M., J.D.J.S., M.J., and A.S.; Visualisation, R.H.P, H.M. and A.S.; Analysis and Software, R.H.P., H.M., and A.S.; Resources, A.A.K. and K.R.S.; Administration, A.A.K., A.S., and K.R.S.; Funding acquisition, A.A.K. and K.R.S.; Writing – original draft, R.H.P., G.M., K.R.S., and A.S.; Writing – review and editing, R.H.P., H.M., G.M., J.D.J.S., M.J., K.R.S., A.S., and A.A.K.

## Acknowledgements

A.A.K. is supported by NHMRC Ideas Grant (2012883) and Australian Infectious Diseases Research Centre (AID) Seed Grant. A.S. was supported by NHMRC Ideas Grant (2021272) and ARC Future Fellowship (FT230100465). The authors acknowledge the facilities and the scientific and technical assistance of the Microscopy Australia Facility at the Centre for Microscopy and Microanalysis (CMM) of The University of Queensland. We thank the Queensland Health Forensic and Scientific Services, Queensland Department of Health, and QIMR Berghofer Medical Research Institute for providing SARS-CoV-2 isolates. Next-generation sequencing was performed by ACE Sequencing (SCMB, UQ). Purchasing of NGS reagents was supported by Illumina COVID Matched Funding Scheme. We are grateful for the gift of C2/E2 SARS-CoV-2 nucleocapsid nanobody from Ariel Isaacs (SCMB, UQ). This work utilised the Australian Galaxy service (https://usegalaxy.org.au/).

