## Supplementary Figures 1-2 for "Omicron evolution drives increased ciliated cell tropism and dysfunction in nasal epithelia"

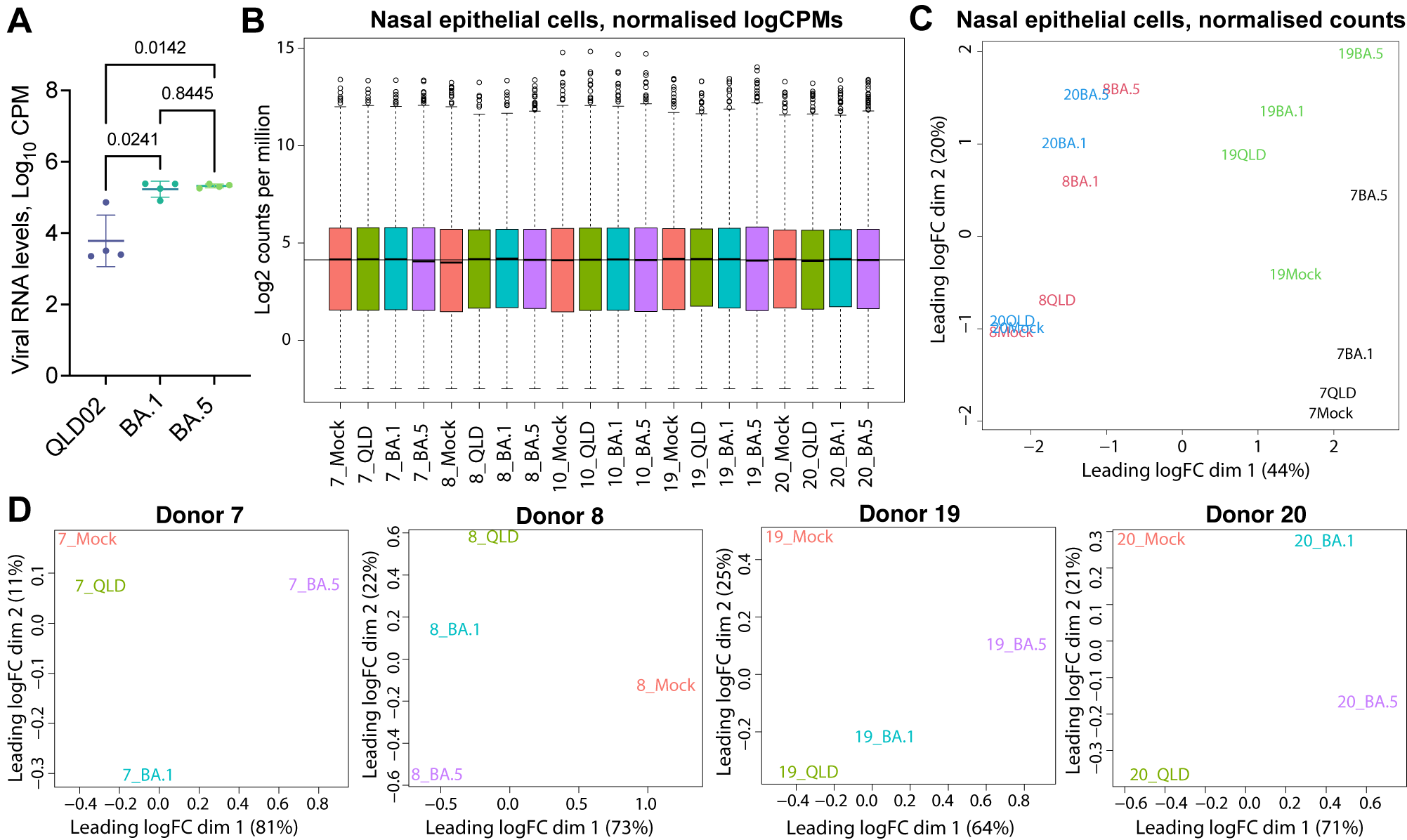

**Supplementary Figure 1. RNA-Seq analysis of primary human nasal epithelial cells after infection with SARS-CoV-2 variants QLD02, BA.1, and BA.5.** (A) Viral RNA levels in human primary nasal epithelial cells (NECs) at 48 hours after infection with QLD02, BA.1, and BA.5 SARS-CoV-2 variants. RNA levels were determined by RNA-Seq followed counting of the reads mapped to the SARS-CoV-2 reference genomes. Each data point represents an independent cell culture derived from a different donor (n = 4). Bars indicate mean values and range. Statistical significance determined using the Kruskal-Wallis test with Dunn’s multiple comparisons. (B) Box plot indicating library sizes and read counts distribution within the RNA-Seq libraries generated from NECs infected with SARS-CoV-2 variants QLD02 (ancestral strain), Omicron BA.1, Omicron BA.5, or mock-infected controls. Libraries were prepared from cells obtained from four independent donors. (C) Global transcriptional distance across all samples used int the RNA-Seq experiment. Multidimensional scaling (MDS) plot of all RNA-Seq libraries, encompassing four donors and four infection conditions. Each point represents a sample, coloured by donor. (D) Donor-stratified MDS plots demonstrating sample clustering according to infection condition (QLD02, BA.1, BA.5, or mock) for each donor.

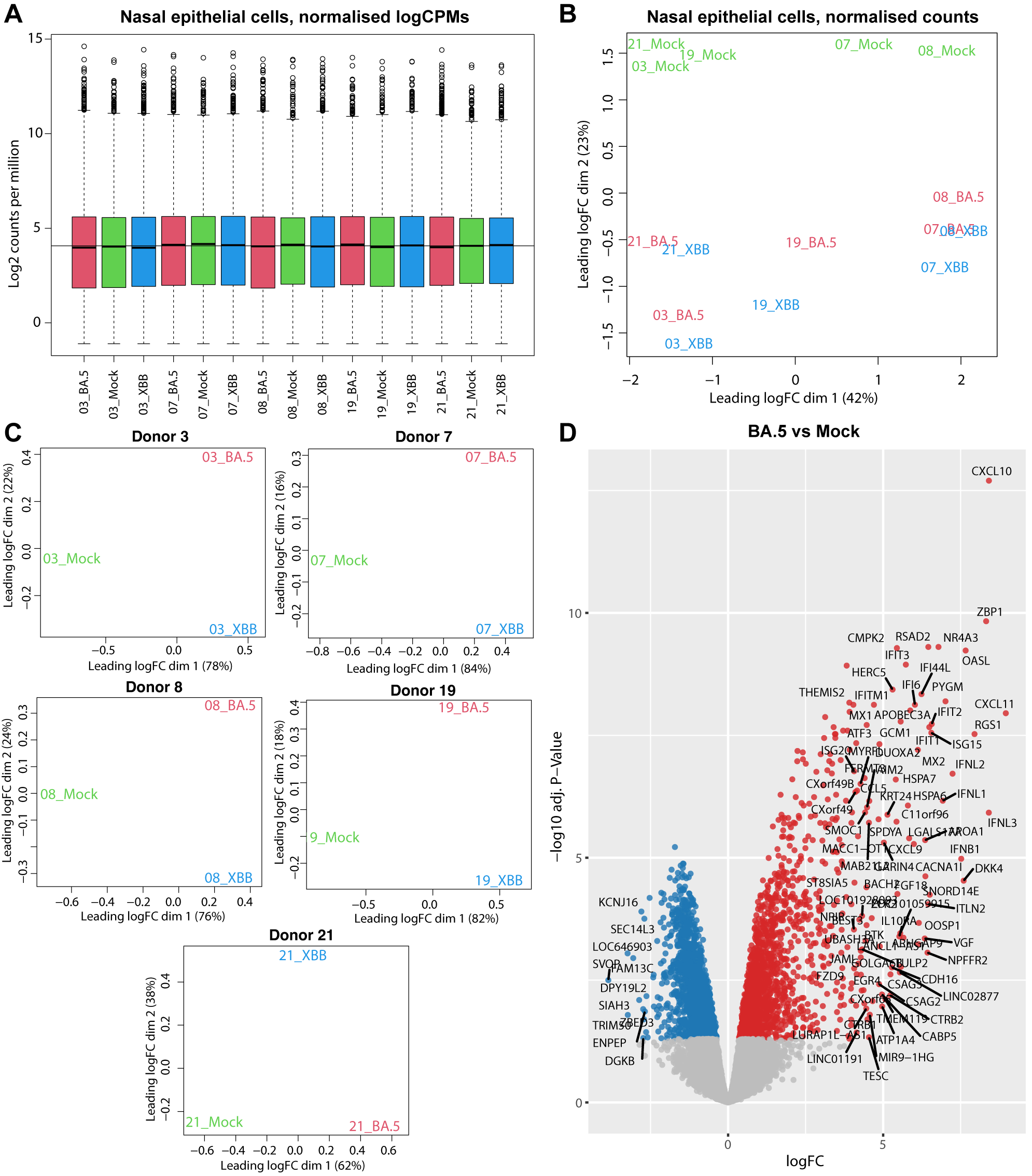

**Supplementary Figure 2. RNA-Seq analysis of human NECs after infection with SARS-CoV-2 variants BA.5 and XBB.** **(A)** Box plot indicating library sizes and read counts distribution within the RNA-Seq libraries generated from human NECs infected with SARS-CoV-2 variants Omicron BA.5, Omicron XBB and mock-infected cells. Libraries were prepared from cells obtained from five independent donors. **(B)** Global transcriptional distance between the samples used in the RNA-Seq experiment. Multidimensional scaling (MDS) demonstrates close clustering of the samples infected with both viruses, with separation from mock-infected samples. Each point represents a sample, coloured by donor. **(C)** Donor-stratified MDS plots demonstrating separation of the infected samples from mock in dimension one, and separation of BA-infected samples from XBB-infected samples in dimension 2. (D) Differentially expressed genes in human NECs infected with SARS-CoV-2 Omicron BA.5 in comparison to mock-infected cells. Data is representative from 5 individual donors. Significantly (FDR-corrected *p* value < 0.05, logFC>1) up-regulated genes are shown in red and down-regulated genes are shown in blue. Top-most significant differentially expressed genes are labelled.
